# PTPN14 Degradation by High-Risk Human Papillomavirus E7 Limits Keratinocyte Differentiation and Contributes to HPV-Mediated Oncogenesis

**DOI:** 10.1101/471045

**Authors:** Joshua Hatterschide, Amelia E. Bohidar, Miranda Grace, Tara J. Nulton, Brad Windle, Iain M. Morgan, Karl Munger, Elizabeth A. White

## Abstract

High-risk human papillomavirus (HPV) E7 proteins enable oncogenic transformation of HPV-infected cells by inactivating host cellular proteins. High-risk but not low-risk HPV E7 target PTPN14 for proteolytic degradation, suggesting that PTPN14 degradation may be related to their oncogenic activity. HPV infects human keratinocytes but the role of PTPN14 in keratinocytes and the consequences of PTPN14 degradation are unknown. Using an HPV16 E7 variant that can inactivate RB1 but cannot degrade PTPN14 we found that high-risk HPV E7-mediated PTPN14 degradation impairs keratinocyte differentiation. Deletion of *PTPN14* from primary human keratinocytes decreased keratinocyte differentiation gene expression. Related to oncogenic transformation, both HPV16 E7-mediated PTPN14 degradation and *PTPN14* deletion promoted keratinocyte survival following detachment from a substrate. PTPN14 degradation contributed to high-risk HPV E6/E7-mediated immortalization of primary keratinocytes and HPV-positive but not HPV-negative cancers exhibit a gene expression signature consistent with PTPN14 inactivation. We find that PTPN14 degradation impairs keratinocyte differentiation and propose that this contributes to high-risk HPV E7-mediated oncogenic activity independent of RB1 inactivation.

**Significance Statement:** Human papillomaviruses uncouple proliferation from differentiation in order to enable virus replication in epithelial cells. HPV E7 proteins are well established to promote proliferation by binding to and inactivating retinoblastoma family proteins and other cell cycle inhibitors. However, mechanisms by which high-risk HPV oncoproteins inhibit differentiation have not been defined. This paper identifies the first mechanism by which high-risk HPV E7 inhibit keratinocyte differentiation. The inhibition of differentiation requires degradation of the cellular protein PTPN14 by high-risk HPV E7 and this degradation is related to the ability of high-risk HPV oncoproteins to immortalize keratinocytes and to cause cancer.

## Introduction

Human papillomaviruses (HPVs) are non-enveloped, double-stranded DNA viruses that infect and replicate in the stratified squamous epithelium. HPV initially infects keratinocytes in the basal, proliferative layer of the epidermis, and subsequent steps in the HPV replicative cycle including viral genome amplification, encapsidation, and egress are dependent on keratinocyte differentiation (1–3). However, HPV genome amplification also requires components of the cellular machinery for DNA replication that are not expressed in differentiating cells. Thus, productive HPV infection must uncouple proliferation and differentiation in the epithelium. Infection with one of the 13-15 ‘high-risk’ HPV causes nearly all cervical cancer, some other anogenital cancer, and an increasing proportion of HPV-positive head and neck squamous cell carcinomas (HNSCC) (4–6). In total, HPV infection causes ~5% of cancers worldwide.

The high-risk HPV E7 oncoprotein is able to immortalize human keratinocytes and the efficiency of immortalization is increased by high-risk HPV E6 (7–9). A well-characterized activity of many HPV E7 is to bind and inactivate the retinoblastoma tumor suppressor (RB1) via the LxCxE motif present in E7 conserved region 2 (CR2) (10–12). In addition, HPV16 E7 can direct the proteasome-mediated degradation of RB1 (13–16). RB1 inactivation releases the inhibition of E2F transcription factors, thus allowing cell cycle progression and acting as a major driver of proliferation. HPV E7 also promote proliferation by inhibiting the CDK inhibitors p21^WAF1/CIP1^ and p27^KIP1^ (17–19). In addition to promoting proliferation, transcriptional studies indicate that human cells harboring high-risk HPV genomes express lower levels of differentiation marker genes and that both high-risk HPV E6 and E7 likely contribute to this repression (20–26). However, a mechanism by which high-risk HPV E6 and/or E7 inhibit differentiation has not been defined.

RB1 binding by HPV E7 is necessary but insufficient for immortalization and transformation, and several observations highlight the need for other contributors to transformation. First, in multiple assays, the oncogenic activity of high-risk HPV E7 is disrupted by mutations in regions that do not include the LxCxE motif (27–31). Second, low-risk HPV E7 bind RB1 but do not have activity in transformation assays and other E7 such as HPV1 E7 bind RB1 with high affinity but do not transform (32–34). Finally, bovine papillomavirus (BPV) E7 does not bind to RB1, but in some assays it is required for BPV-mediated transformation (30, 35–37). The idea that RB1 inactivation is insufficient for transformation is additionally supported by studies in mouse models of cervical cancer (38, 39). Overall, updates to the model of transformation by HPV E6 and E7 have been suggested (40) and additional binding partners of E7 have been proposed to mediate transformation independent of RB1 binding (41–43). However not all of these interactions are conserved among the high-risk HPV E7.

The E3 ubiquitin ligase UBR4 is a conserved interactor of diverse papillomavirus E7 (44). UBR4 is required by both HPV16 E7 and BPV E7 for RB1-independent transformation but for some years the reason for this requirement was unknown (45, 46). Recently we discovered that the cellular protein PTPN14 binds to HPV E7 proteins from diverse HPV genotypes and that high-risk HPV E7 use UBR4 to direct PTPN14 for proteasome-mediated degradation. Although low-risk HPV E7 also bind UBR4, only high-risk HPV E7 mediate PTPN14 degradation, and E7 binding to PTPN14 and to UBR4 does not require interaction with RB1 (44, 47).

PTPN14 is a non-receptor protein tyrosine phosphatase that is evolutionarily conserved as a regulator of developmental signaling from *Drosophila* to humans, however phenotypes associated with PTPN14 loss vary (48–52). Hereditary variations in human *PTPN14* are associated with developmental disorders including dysregulated angiogenesis, improper lymphatic development and improper choanal development (48, 51). Mutations in human cancer have implicated PTPN14 as a putative tumor suppressor (53–56). *PTPN14* is mutated in cancers such as colorectal cancer and basal cell carcinoma and in both cancer types mutations occur along the length of the gene (54, 57). Several potential substrates for dephosphorylation by PTPN14 are related to cell growth control (53, 56, 58). PTPN14 also has phosphatase independent activities such as the ability to regulate Hippo signaling through direct interaction with YAP1 or with its upstream regulators LATS1/2 (55, 59–61). These interactions are mediated through central PPxY motifs in PTPN14.

Based upon the observations that the ability of E7 to degrade PTPN14 correlates with E7 oncogenic activity, that the regions of high-risk HPV E7 required for PTPN14 degradation are the same as those that confer RB1-independent transforming activity, and that *PTPN14* is a putative tumor suppressor, we hypothesized that PTPN14 degradation could be required for high-risk HPV E7-mediated oncogenic transformation. The biological activities of PTPN14 in keratinocytes have not been studied, and the molecular consequences of PTPN14 degradation by high-risk HPV E7 have not been defined. Here we report that PTPN14 loss impaired the differentiation program in human keratinocytes and that HPV16 E7 could inhibit the expression of differentiation marker genes in response to stimulus. This inhibition was dependent upon HPV16 E7’s ability to degrade PTPN14 and was retained in the absence of RB1 binding. Moreover, the ability of E7 to degrade PTPN14 contributed to the immortalization of primary human keratinocytes by HPV16 E6 and E7. Repression of differentiation is a potentially oncogenic event and we found that repression of keratinocyte differentiation describes the major gene expression differences between HPV+ and HPV-HNSCC. Taken together, our results suggest that high-risk E7 mediated PTPN14 degradation impairs keratinocyte differentiation. This is an RB1-independent, and potentially oncogenic, activity of high-risk HPV E7.

## Results

### The HPV16 E7 E10K variant is impaired in PTPN14 degradation but binds RB1 and promotes E2F target gene expression

PTPN14 degradation by high-risk HPV E7 requires the E3 ubiquitin ligase UBR4, which interacts with the N-terminus of E7. PTPN14 binding maps broadly to the E7 C-terminus (Figure 1A). The recent identification of HPV16 E7 variants from over 5000 patient samples (62) prompted us to test whether an N-terminal variant might be impaired in the ability to degrade PTPN14. One variant, HPV16 E7 E10K (glutamic acid to lysine change at amino acid 10), is altered in the region that is required for binding to UBR4. To assess the biological activities of this E7 variant we used hTert-immortalized human foreskin keratinocytes (N/Tert-1) (63) to establish cell lines that stably express Flag and HA epitope tagged versions of the prototypical HPV16 E7 (WT), the HPV16 E7 E10K variant, HPV16 E7 Δ21-24, or an empty vector control. The Δ21-24 deletion eliminates the LxCxE motif that is required for E7 to bind to RB1 (12). HPV16 E7 cells exhibited reduced PTPN14 protein levels and binding to RB1 was not required for this effect (Figure 1B). However, HPV16 E7 E10K did not promote the reduction in steady-state PTPN14 protein levels. UBR4 did not co-immunoprecipitate with the HPV16 E7 E10K variant (Figure 1B), suggesting that this variant cannot target PTPN14 for degradation because it is deficient in binding to the required E3 ubiquitin ligase. HPV16 E7 E10K was comparable to HPV16 E7 WT in its ability to bind RB1 (Figure 1B). In primary human foreskin keratinocytes (HFK) stably transduced with the same retroviral vectors, both HPV16 E7 WT and HPV16 E7 E10K could induce the expression of E2F target genes *CCNE1* and *MCM2* (Figure 1C). This supported the notion that UBR4 binding and PTPN14 degradation by HPV16 E7 is independent of RB1 binding and established HPV16 E7 E10K as a variant that is RB1 binding/degradation competent but cannot degrade PTPN14.

**Figure 1.**
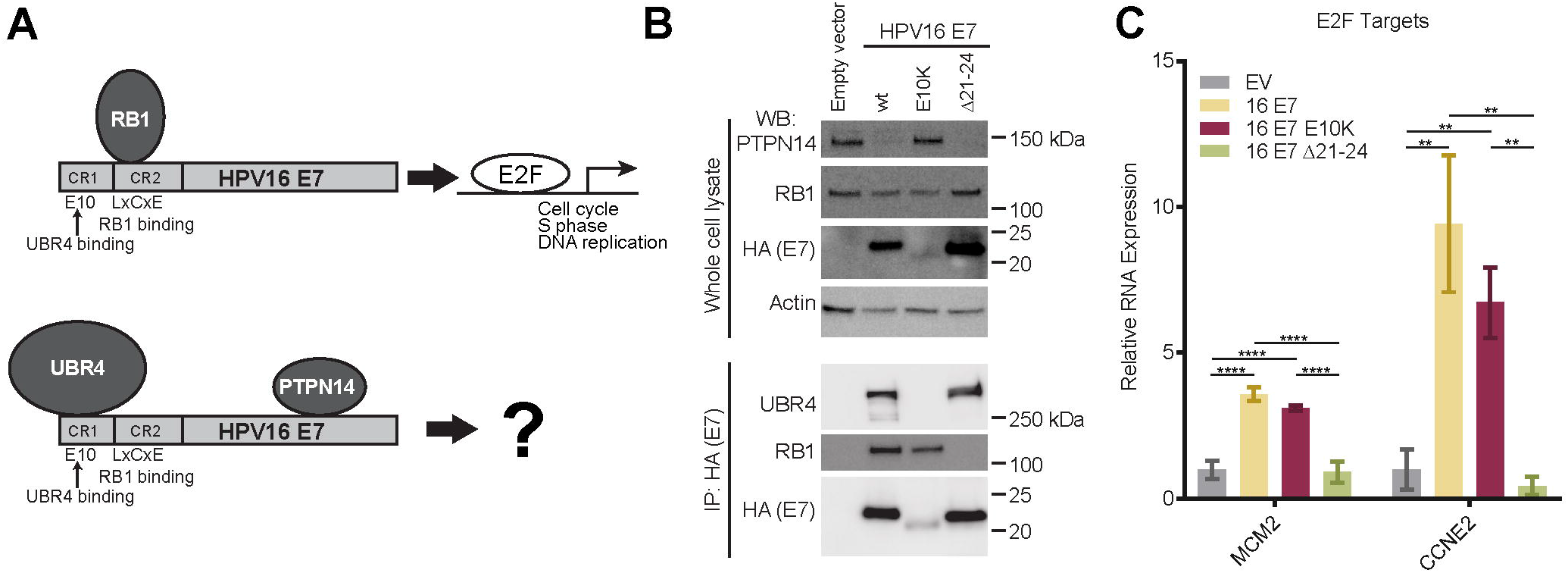
The HPV16 E10K variant is impaired in PTPN14 degradation but binds RB1 and promotes the expression of E2F-regulated genes. (A) Schematic of protein complexes including HPV E7/RB1 and HPV E7/PTPN14/UBR4. (B) N/Tert-1 keratinocytes were transduced with control and HPV16 E7 retroviruses. Total cell lysates were analyzed by SDS-PAGE/Western blotting and probed with antibodies to PTPN14, RB1, HA, and actin (top). HPV16 E7-FlagHA was immunoprecipitated with anti-HA from N/Tert lysates and co-immunoprecipitation of UBR4 and RB1 was assessed by SDS-PAGE/Western blotting (bottom). (C) qRT-PCR for E2F-regulated genes in primary HFK transduced with control and HPV16 E7 retroviruses. Bar graphs display the mean ± standard deviation of 2 (16E7 Δ21-24) or 3 (EV, 16E7 WT, and 16E7 E10K) independent experiments. Statistical significance was determined by ANOVA followed by multiple t-tests with the Holm-Šídák family-wise error rate correction (** = *p* <0.01; *** = *p* < 0.001; **** = *p* < 0.0001).

### HPV16 E7 degrades PTPN14 to inhibit keratinocyte differentiation

To determine whether HPV16 E7 has effects on cellular gene expression that are dependent on its ability to degrade PTPN14, we performed an unbiased analysis of gene expression in keratinocytes expressing HPV16 E7 variants. Duplicate or triplicate primary HFK cell populations were established by transduction with retroviral vectors encoding HPV16 E7 WT, HPV16 E7 E10K, and HPV16 E7 Δ21-24 and selected with puromycin. Total RNA was isolated from independent cell populations then polyA selected RNA was subjected to RNA-seq. As predicted by our initial validation of the E10K variant, HPV16 E7 E10K behaved like HPV16 E7 WT with respect to the upregulation of DNA replication genes and had a comparable effect on genes related to RB1 binding (Supplemental Table 1 and Supplemental Figure 1).

Next, we assessed the differences between HPV16 E7 WT and HPV16 E7 E10K. Seventy-five genes were differentially regulated in HPV16 E7 E10K cells compared to HPV16 E7 WT cells with fold change ≥1.5 and adjusted *p*-value ≤0.05. Approximately half of these differentially-regulated genes were repressed more by HPV16 E7 WT than by HPV16 E7 E10K. Gene ontology (GO) enrichment analysis showed that the genes repressed by HPV16 E7 dependent on its ability to degrade PTPN14 were described by developmental GO terms that are related to the keratinocyte differentiation program (Figure 2A). These included epithelial cell differentiation, cornification, keratinocyte differentiation, epidermis development, epidermal cell differentiation, and keratinization. Many of the individual genes that were repressed by HPV16 E7 but not by HPV16 E7 E10K are differentiation markers (Figure 2B, y-axis). In contrast, genes that were activated by HPV16 E7 dependent on its ability to degrade PTPN14 were not significantly enriched for any GO terms (Supplemental Figure 2A).

**Figure 2.**
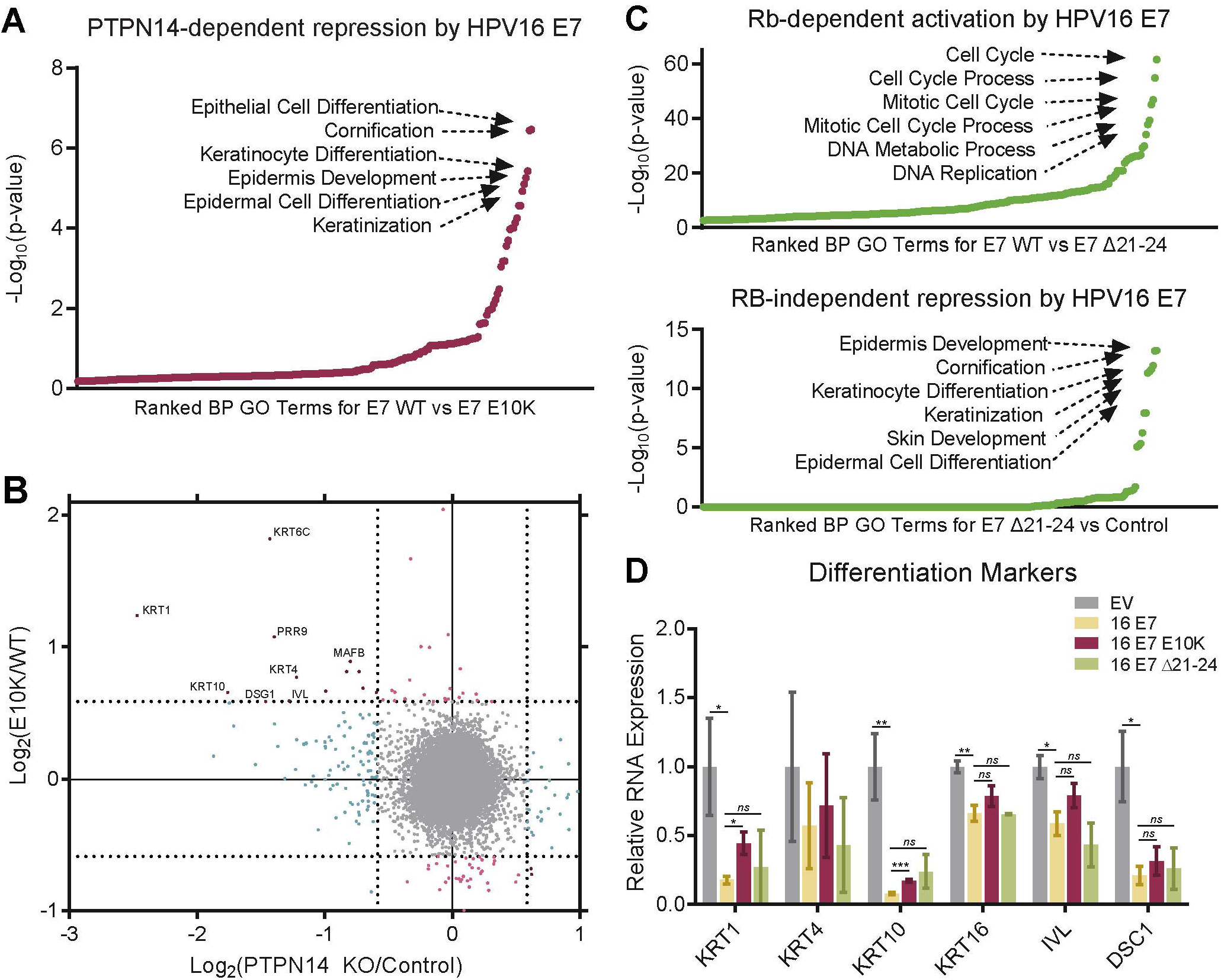
HPV16 E7-mediated degradation of PTPN14 inhibits keratinocyte differentiation. Primary HFK were transduced with retroviruses encoding HPV16 E7, HPV16 E7 E10K, HPV16 E7 Δ21-24, or an empty vector control. PolyA selected RNA was analyzed by RNA-seq. (A) GO enrichment analysis of genes with ≥1.5 fold lower expression in HPV16 E7 WT cells relative to HPV16 E7 E10K cells and *p*-value ≤0.05. (B) Scatter plot of log_2_(fold-change) in gene expression compares the gene expression changes of HPV16 E7 E10K relative to HPV16 E7 WT to those of PTPN14 KO relative to control. Colors denote whether genes are altered by PTPN14 KO only (blue), by E7 WT more than E7 E10K only (light red), or both (dark red). (C) Same analysis as (A) of (C, Top) genes with ≥1.5 fold higher expression in HPV16 E7 WT than HPV16 E7 Δ21-24 cells, and (C, Bottom) genes ≥1.5 fold lower expression in HPV16 E7 Δ21-24 cells relative to empty vector control cells and *p*-value ≤0.05. (D) Impacts of HPV16 E7 WT, HPV16 E7 E10K, and HPV16 E7 Δ21-24 on gene expression in primary HFK cells were validated by qRT-PCR targeting markers of differentiation. Bar graphs display the mean ± standard deviation of 2 (16E7 Δ21-24) or 3 (EV, 16E7 WT, and 16E7 E10K) independent experiments. Statistical significance was determined by ANOVA followed by multiple t-tests with the Holm-Šídák family-wise error rate correction (* = *p* <0.05; ** = *p* <0.01; *** = *p* < 0.001).

To test whether repression of keratinocyte differentiation was related to RB1 inactivation we used HPV16 E7 Δ21-24 to assess the transcriptional impact of E7 in the absence of RB1 binding. As expected, cell cycle and DNA replication related GO categories were the most significantly enriched categories among genes differentially regulated by HPV16 E7 WT versus HPV16 E7 Δ21-24 (Figure 2C, Top). In contrast, comparing HPV16 E7 Δ21-24 to empty vector control identified the genes that are repressed by HPV16 E7 independent of RB1 binding (Figure 2C, Bottom). GO analysis of these genes identified the same keratinocyte differentiation-related gene sets that were seen in our analysis of PTPN14 degradation dependent effects of HPV16 E7. Furthermore, individual genes repressed by HPV16 E7 Δ21-24 relative to control (Supplemental Figure 2C, y-axis) are similar to those repressed by HPV16 E7 WT dependent on its ability to degrade PTPN14. We concluded that repression of keratinocyte differentiation through the degradation of PTPN14 was independent of RB1 binding. In the absence of RB1 binding HPV16 E7 acted mainly as a repressor but retained a modest ability to promote gene expression (Supplemental Figure 2B).

To better understand the impacts of PTPN14 degradation on gene expression, we examined individual genes significantly lower in HPV16 E7 WT cells than in HPV16 E7 E10K cells. Many of the genes that are repressed by HPV16 E7 WT and HPV16 E7 Δ21-24 but not by HPV16 E7 E10K are described by epidermis development and more specific GO terms (Supplemental Figure 2D, Supplemental Table 2). Although certain genes were not repressed by either HPV16 E7 E10K or HPV16 E7 Δ21-24, these genes were largely related to other biological processes.

To validate the results obtained from RNA-seq, we used qRT-PCR to confirm the altered expression of several genes related to keratinocyte differentiation in our cell lines. Markers of keratinocyte differentiation such as keratin 1 (KRT1), keratin 4 (KRT4), keratin 10 (KRT10), keratin 16 (KRT16), involucrin (IVL), and desmocollin 1 (DSC1) were repressed by HPV16 E7 by 1.5-to 12-fold (Figure 2D). KRT1, KRT4, KRT10, and KRT16 are cytokeratins associated with the suprabasal layers of differentiating keratinocytes. IVL constitutes a major component of the cornified envelope and is expressed at high levels in the upper layers of the epidermis. DSC1 is a component of desmosome complexes associated with keratinization and is expressed at higher levels in the upper spinous layer and granular layer of the epidermis. Comparison of both HPV16 E7 WT and Δ21-24 to the HPV16 E7 E10K variant indicated that the ability of HPV16 E7 to repress expression of these genes was at least partially dependent on its ability to target PTPN14 for degradation.

### The ability of E7 to degrade PTPN14 correlates with its ability to inhibit differentiation and promote survival upon detachment

Next, we wanted to determine whether the ability of HPV E7 to alter differentiation-related gene expression in unstimulated cells correlated with changes following a differentiation stimulus. *In vivo*, detachment from the basement membrane stimulates keratinocyte differentiation, an effect that can be mimicked by growth of cultured cells in suspension (64–66). HPV E7 has been previously shown to protect against cell death following detachment in a UBR4-dependent manner (46, 67). In these experiments we used N/Tert-1 cells engineered to stably express HPV16 E7 WT, HPV16 E7 E10K, HPV6 E7, or an empty vector control. HPV6 is a low-risk HPV encoding an E7 that, like HPV16 E7 E10K, binds PTPN14 but does not direct it for proteasome-mediated degradation. Using immortalized cells in these experiments enabled propagation of sufficient numbers of cells for detachment assays.

N/Tert-1 cells were harvested directly from adherent culture or subjected to growth in suspension for 12h to induce differentiation. KRT16 and IVL RNAs were analyzed by qRT-PCR and were induced by detachment in all of the cell lines tested. Detached empty vector cells expressed 7-to 30-fold more of these transcripts compared to adherent cells and each version of E7 limited the induction of KRT16 and IVL. KRT16 and IVL expression was 2.3- or 3.4-fold lower in N/Tert-HPV16 E7 WT cells compared to the empty vector control and repression of IVL was statistically significant (Figure 3A). The statistical significance of some other comparisons was limited by the fact that there was a wide range of induction of the differentiation markers following detachment. However, the trend was highly reproducible: in three replicate experiments HPV16 E7 WT always repressed differentiation marker gene expression more than HPV16 E7 E10K and HPV6 E7. These results indicate that PTPN14 degradation is required for maximal repression of detachment-induced differentiation by HPV16 E7.

**Figure 3.**
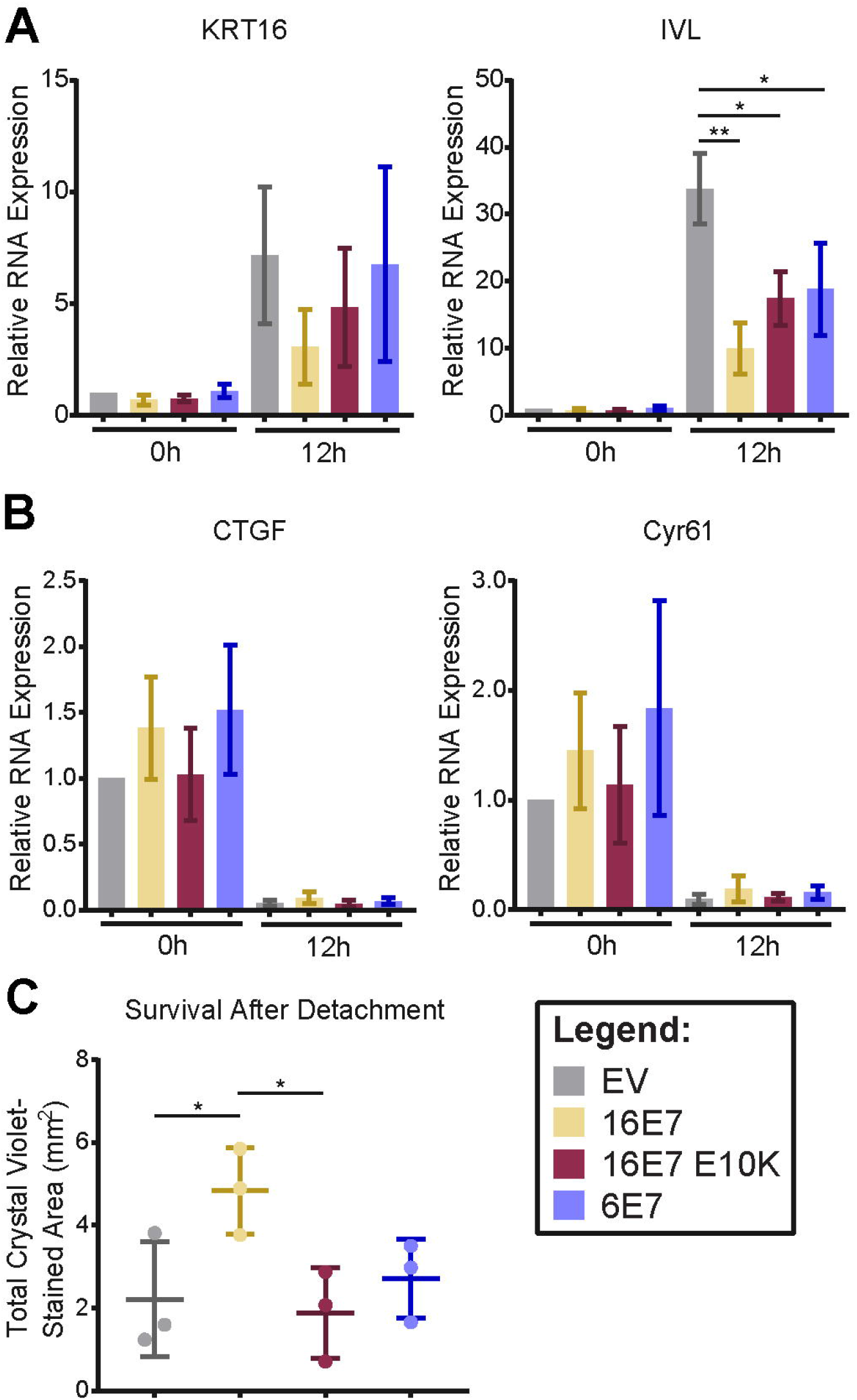
The ability of E7 to degrade PTPN14 correlates with its ability to inhibit differentiation and promote survival upon detachment. N/Tert-1 stably transduced with retroviruses encoding HPV16 E7, HPV16 E7 E10K, HPV6 E7, or an empty vector control were subjected to growth in suspension for 12h and assayed for markers of differentiation, YAP/TEAD targets, and survival after detachment. (A and B) Gene expression changes induced by suspension were assayed by qRT-PCR targeting markers of differentiation: KRT16 and IVL (A), and YAP/TEAD targets: CTGF and CYR61 (B). mRNA expression was calculated relative to GAPDH. Bar graphs display the mean ± standard deviation of 3 independent experiments. (C) Survival after detachment was assayed by replating 1,000 cells from suspension and measuring the surface area covered after 5 days of growth by crystal violet staining. Three independent experiments are displayed along with mean ± standard deviation. Statistical significance was determined by ANOVA followed by multiple t-tests with the Holm-Šídák family-wise error rate correction (* = *p* <0.05; ** = *p* <0.01)

Several explanations could account for the observation that E7 that do not degrade PTPN14 still partially repress differentiation. Each of the E7 tested here bind and inactivate RB1 and it is possible that some inhibition of differentiation is due to the increased proliferation resulting from RB1 inactivation. Another, not mutually exclusive, explanation could be that PTPN14 binding alone is enough to result in some inhibition of differentiation. Our data support this idea, since HPV16 E7 E10K and HPV6 E7 both interact with PTPN14 and they repressed differentiation to similar levels.

In addition to stimulating differentiation, growth in suspension activates the Hippo signaling pathway (68, 69) which represses the transcription of the well characterized YAP/TEAD targets *CTGF* and *CYR61*. PTPN14 knockdown in MCF10A cells has been shown to induce the transcription of *CTGF* and *CYR61* (59). Since PTPN14 has been characterized as a negative regulator of YAP1 and shown to regulate *CTGF* and *CYR61* in other cell types (55, 59–61), we measured these transcripts to determine whether E7 differentially impacts their expression. Compared to vector controls, none of the E7 cell lines exhibited altered expression of these YAP1/TEAD targets before or after detachment (Figure 3B).

To further assess cell viability in the detachment experiment, 1,000 cells were taken from suspension culture, re-plated in coated tissue culture plates, and allowed to grow for 5d. HPV16 E7 protected against cell death following detachment in a manner that was dependent on the ability of E7 to target PTPN14 for degradation (Figure 3C). This is consistent with previous reports demonstrating that HPV16 E7 and BPV E7 require UBR4 to protect cells against cell death triggered following detachment from a substrate (46, 67).

### PTPN14 knockout limits differentiation gene expression in primary human keratinocytes

To test what cellular processes are affected when PTPN14 levels are reduced in human keratinocytes, we performed an unbiased analysis of gene expression in the presence and absence of PTPN14. Primary HFK were transduced with lentiviral vectors encoding SpCas9 plus an sgRNA targeting PTPN14 (sgPTPN14-3) or a nontargeting control sgRNA (sgNT-2), then selected with puromycin to generate control (HFK-control) and PTPN14-deleted (HFK-PTPN14 KO) cell lines (Figure 4A). Total RNA was isolated from two or three independent isolates of HFK-control and HFK-PTPN14 KO, then polyA selected RNA was subjected to RNA-seq. In cells that did not express *PTPN14*, 141 genes were differentially regulated with fold change ≥1.5 and adjusted *p*-value ≤0.05. Of these 29 genes were up-regulated and 112 were down-regulated in the absence of PTPN14 (Figure 4B and Supplemental Table 3). Thus, PTPN14 appeared to act largely to promote, rather than to repress, gene expression. As in the analysis of the HPV16 E7 variants, keratinocyte differentiation-related GO terms were downregulated in PTPN14 KO cells (Figure 4C, Figure 2A and C). More than half of the down-regulated genes were in epidermis development-related or other developmental process-related GO categories (Figure 4D). There was not a corresponding enrichment in differentiation-related GO terms among the up-regulated genes; however, one GO category, inflammatory response, was significantly enriched in this analysis (Supplemental Figure 3).

**Figure 4.**
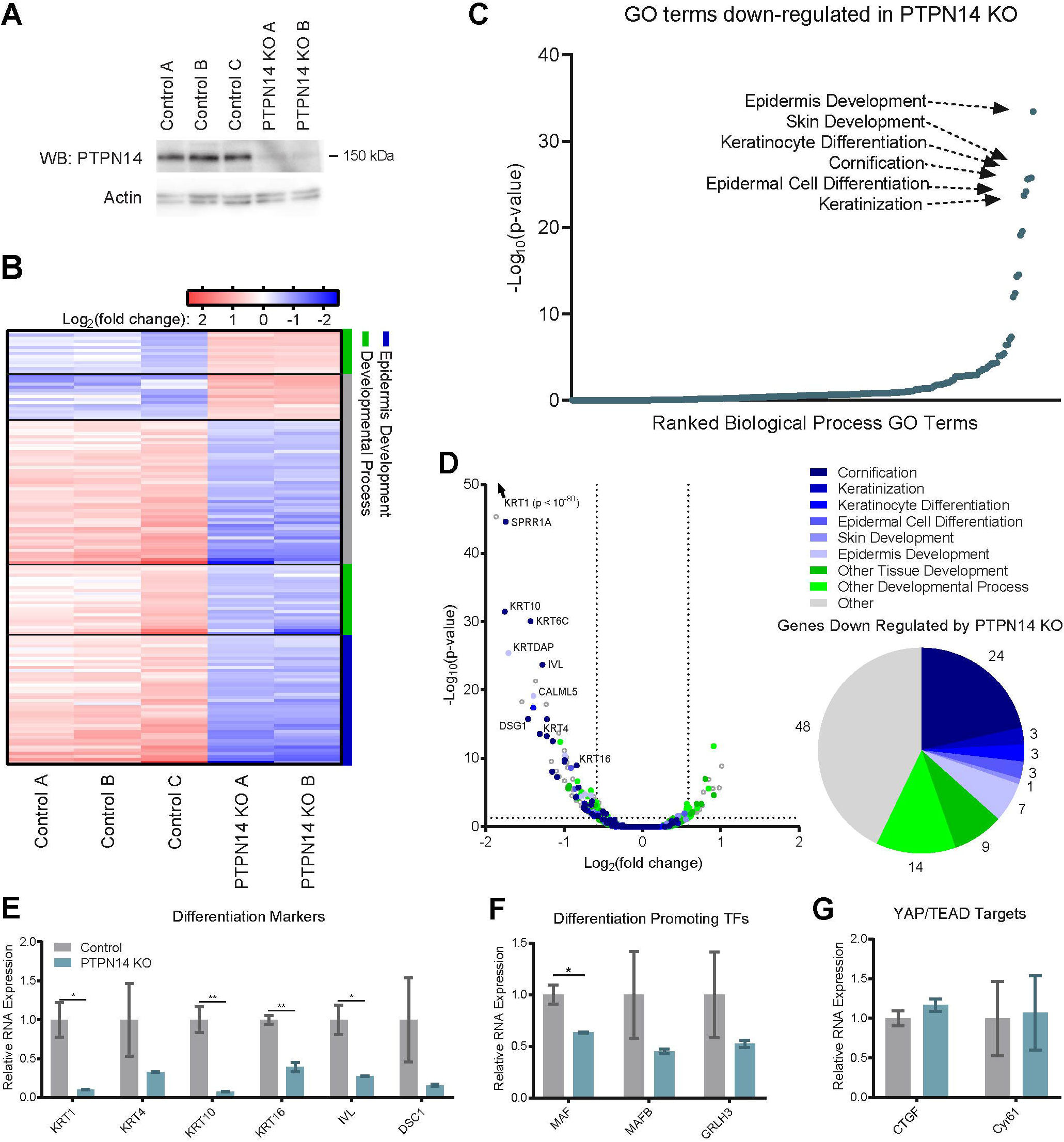
PTPN14 depletion impairs differentiation-related gene expression in primary human keratinocytes. Primary HFK were transduced with LentiCRISPRv2 lentiviral vectors encoding SpCas9 and non-targeting or PTPN14-directed sgRNAs and analyzed for changes in gene expression. (A) Cell lysates were subjected to SDS-PAGE/Western analysis and probed with anti-PTPN14 and anti-actin antibodies. (B) PolyA selected RNA was analyzed by RNA-seq. Genes differentially expressed by >1.5 fold with *p*-value ≤0.05 are displayed in the heat map. Color coding on the right side denotes whether genes are related to epidermis development (blue), other developmental processes (green), or neither (gray). (C) GO enrichment analysis of genes downregulated in HFK-PTPN14 KO compared to HFK-control. (D) Volcano plot of gene expression changes in HFK-control vs HFK-PTPN14 KO. Dots colored by GO terms. Pie chart displays the fraction of genes down regulated in the absence of PTPN14 that fall into enriched GO Terms. (E, F, and G) Transcript abundance for selected genes in HFK-control and HFK-PTPN14 KO was measured by qRT-PCR detecting differentiation markers (E), differentiation promoting transcription factors (F), and YAP/TEAD targets (G). Bar graphs display the mean ± standard deviation of 2 or 3 independent experiments. Statistical significance was determined by Welch’s t-tests (* = *p* <0.05; ** = *p* <0.01).

We hypothesized that individual genes might be similarly regulated by PTPN14 KO and E7-mediated PTPN14 degradation. Indeed, the genes that were both downregulated by PTPN14 loss and downregulated by HPV16 E7 WT in a PTPN14 degradation-dependent manner are involved in keratinocyte differentiation (Figure 2B). Furthermore, gene expression changes induced by HPV16 E7 Δ21-24 are positively correlated with those resulting from PTPN14 KO (Supplemental Figure 2C). Taken together, gene expression analysis of HFK-PTPN14 KO and cells expressing HPV16 E7 variants is consistent with degradation of PTPN14 by HPV16 E7 acting to inhibit keratinocyte differentiation. Our data suggest that PTPN14 degradation mediates the predominant RB-independent effect of HPV16 E7 on gene expression.

We selected a subset of genes for validation by qRT-PCR. In agreement with the RNA-seq results, markers of keratinocyte differentiation such as *KRT1, KRT4, KRT10, KRT16, IVL*, and *DSC1* were expressed at 3- to 12-fold lower levels in the absence of PTPN14 (Figure 4E). Transcription factors (TFs) such as MAF, MAFB, and GRLH3 that are transcriptionally regulated during progression of the keratinocyte differentiation program (70–74) exhibited lower expression in the absence of PTPN14 (Figure 4F). Unlike the published effects in other cell types, we found that PTPN14 loss did not impact the expression of the well-characterized YAP/TEAD targets *CTGF* and *CYR61* (Figure 4G). These data support the idea that PTPN14 loss impairs the regulation of keratinocyte differentiation but does not affect expression of canonical Hippo regulated genes in primary HFK.

### PTPN14 contributes to the upregulation of differentiation markers upon detachment

Having determined that PTPN14 loss reduces the basal expression of keratinocyte differentiation-related genes, we next tested whether PTPN14 loss alters the cellular response to a differentiation stimulus. We used CRISPR-Cas9 gene editing in N/Tert-1 cells to engineer control (N/Tert-mock) or PTPN14-deleted (N/Tert-PTPN14 KO) pooled stable cell lines. Again, we stimulated these cells to differentiate through growth in low adherence plates for 12h. Consistent with the effect in primary cells, PTPN14 KO reduced the expression of *KRT16* and *IVL* in adherent cells. We further found that PTPN14 loss also impaired the expression of *KRT16* and *IVL* upon the induction of differentiation (Figure 5A), mirroring the results observed in our N/Tert-E7 cells. As we observed in the N/Tert-E7 cells as well as the primary HFK-PTPN14 KO cells, N/Tert-PTPN14 KO cells did not express significantly more *CTGF* or *CYR61* than mock controls in either the adherent condition or following growth in suspension (Figure 5B).

**Figure 5.**
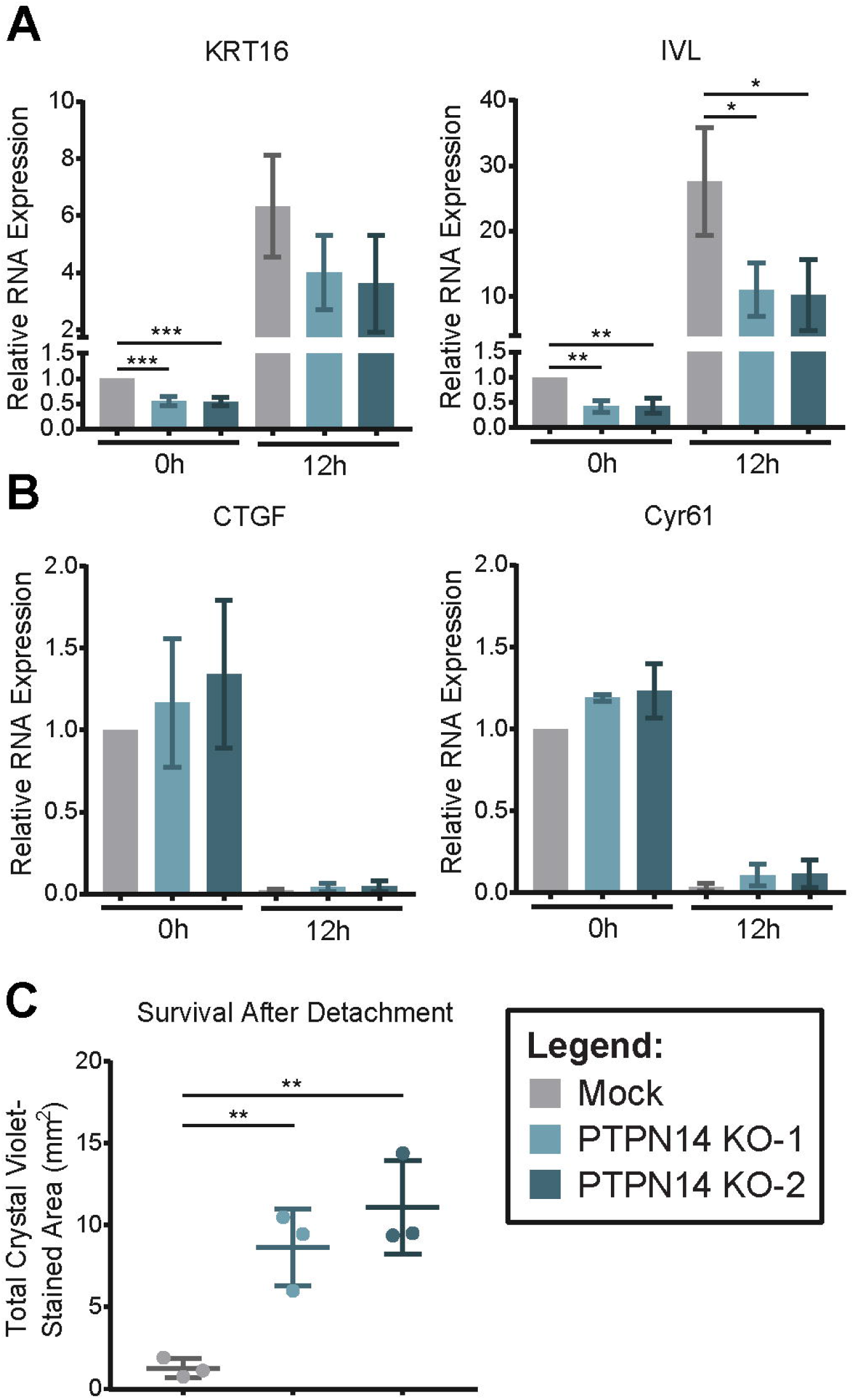
PTPN14 loss reduces the expression of differentiation markers after detachment. Adherent N/Tert-mock and -PTPN14 KO cells were detached by trypsinization and re-plated in ultra-low adherence plates before harvesting 12h post detachment, then assayed for markers of differentiation, YAP/TEAD targets, and survival after detachment. (A and B) Gene expression changes induced by suspension were assayed by qRT-PCR targeting markers of differentiation: KRT16 and IVL (A), and YAP/TEAD targets: CTGF and CYR61 (B). mRNA expression was calculated relative to GAPDH. Bar graphs display the mean ± standard deviation of 3 independent experiments. (C) Survival after detachment was assessed by re-plating 1,000 cells from suspension, allowing cells to grow for 5 days, and measuring total viable cell area by crystal violet staining. Three independent experiments are displayed along with mean ± standard deviation. Statistical significance was determined by ANOVA followed by multiple t-tests with the Holm-Šídák family-wise error rate correction (* = *p* <0.05; ** = *p* <0.01; *** = *p* <0.001)

Finally, we used cell growth after re-plating as a measure of viability after detachment. The N/Tert-PTPN14 KO cells exhibited improved survival and colony formation after detachment compared to control cells (Figure 5C). This is consistent with the result that HPV E7 expression improved survival after suspension in a PTPN14 degradation-dependent manner (Figure 3C) and indicates that loss of PTPN14 is sufficient to improve survival of keratinocytes after detachment.

### PTPN14 degradation contributes to E6/E7 immortalization of primary human keratinocytes

Coexpression of HPV16 E6 and E7 can efficiently immortalize primary keratinocytes in cell culture. To determine whether PTPN14 degradation is required for immortalization by HPV16 oncoproteins, primary HFK were transduced with pairs of HPV E6/E7-encoding retroviruses, selected with puromycin and blasticidin, and monitored for cell growth over the next 17 passages - equivalent to 75 days for WT HPV16 E6/E7 cells (Figure 6).

**Figure 6.**
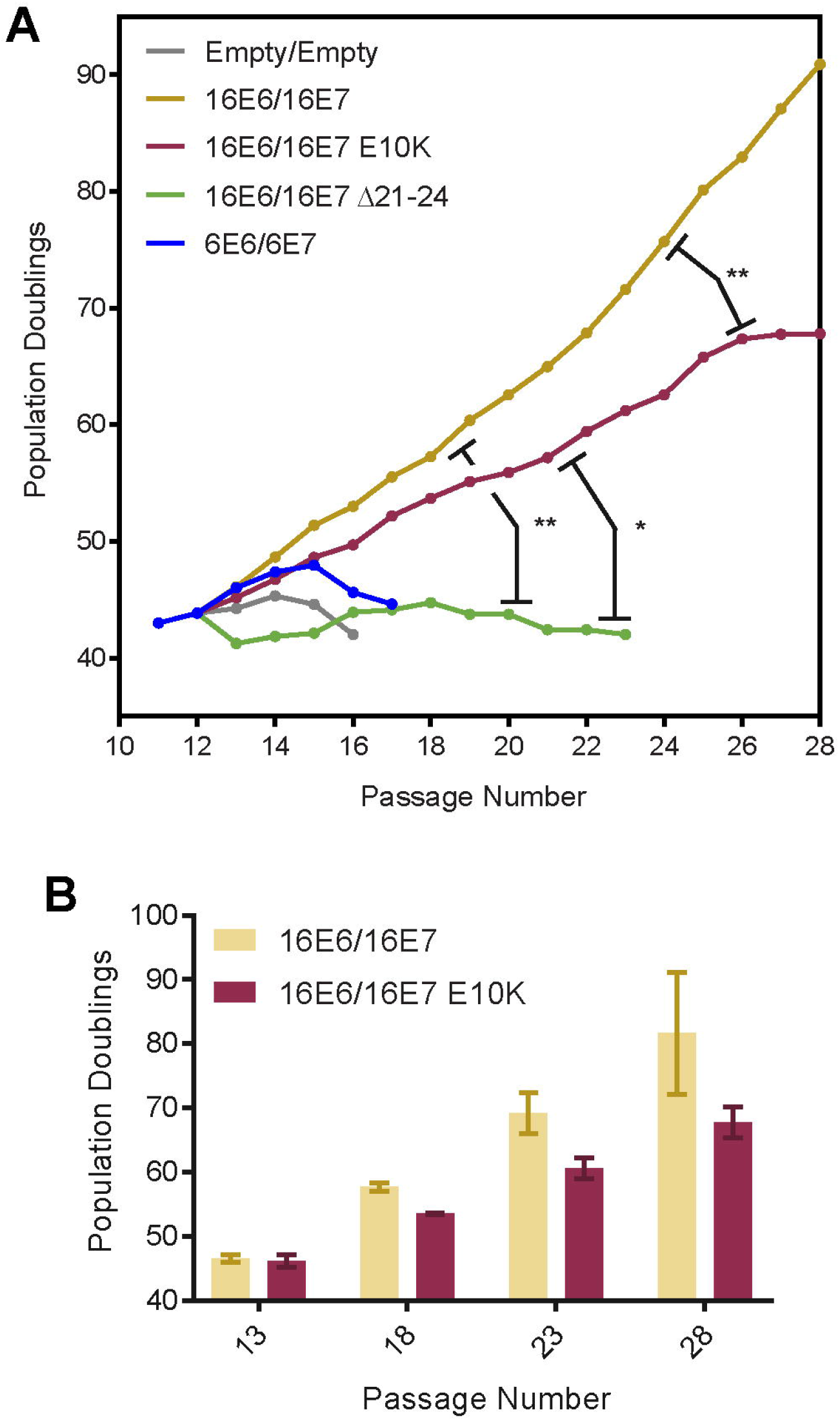
PTPN14 degradation contributes to E6/E7 immortalization of primary human keratinocytes. Primary HFK cells were transduced with pairs of retroviruses encoding various E6 and E7 and passaged for up to 75 days. (A) Growth curves from representative immortalization experiment. Population doublings were calculated based upon the number of cells harvested at each passage. Statistical significance was determined from 3 independent experiments by repeated measures two-way ANOVA. Displayed *p*-values represent the column (cell line) factor (* = p < 0.05; ** = p < 0.01). (B) Bar chart shows mean ± standard deviation of selected passages from growth curves in (A).

Primary HFK transduced with HPV6 E6/E7 or with empty vector controls rapidly senesced, while cells transduced with HPV16 E6/E7 were immortalized in 3/3 replicate experiments. The cells transduced with HPV16 E6/E7 Δ21-24 retroviruses were severely growth impaired and were not immortalized but exhibited a small degree of lifespan extension, perhaps due to sporadic epigenetic inactivation of RB1. Cells transduced with HPV16 E6/E7 E10K expressing vectors retained some proliferative capacity, but their growth was reproducibly impaired compared to that of HPV16 E6/E7 WT cells. We hypothesize that these cells are not fully immortalized and that both RB1 inactivation and PTPN14 degradation are required for immortalization of primary HFK by HPV E6 and E7.

### Keratinocyte differentiation gene expression describes the major differences between HPV+ and HPV-HNSCC

The changes in differentiation-related gene expression in HPV E7-expressing cell lines appeared to be dependent on the ability of E7 to degrade PTPN14 and to reflect the same changes that result from PTPN14 loss in primary HFK. The ability of E7 to degrade PTPN14 also correlates with its ability to immortalize primary HFK. We wished to determine whether E7- or PTPN14-dependent changes in differentiation-related gene expression are reflected in HPV-associated cancers. Using RNA-seq data from the Cancer Genome Atlas (TCGA) we examined gene expression signatures in 508 HNSCC samples, 60 of which are HPV-positive and 448 HPV-negative (75–77). Genes that were differentially expressed by 3-fold or more in HPV-positive vs. HPV-negative samples were selected for further analysis.

Strikingly, the most enriched GO terms among genes downregulated in HPV+ cancers relative to HPV- cancers were epidermis development, keratinocyte differentiation, and epidermal cell differentiation (Figure 7A). As in the PTPN14 knockout cells and in the presence of HPV16 E7, downregulated genes reflected a keratinocyte differentiation signature. Furthermore, many of the other highly enriched GO terms were related to more general developmental processes. In total, epidermis development and other developmental processes accounted for about one-third of the differentially regulated genes in HPV-positive vs. HPV-negative HNSCC. In contrast GO enrichment identified no clear gene sets enriched among genes upregulated in HPV-positive compared to HPV-negative HNSCC (Supplemental Figure 4). The downregulation of differentiation-related genes in HPV-positive relative to HPV-negative cancers is consistent with the changes in gene expression induced by the high-risk HPV E7-mediated degradation of PTPN14.

**Figure 7.**
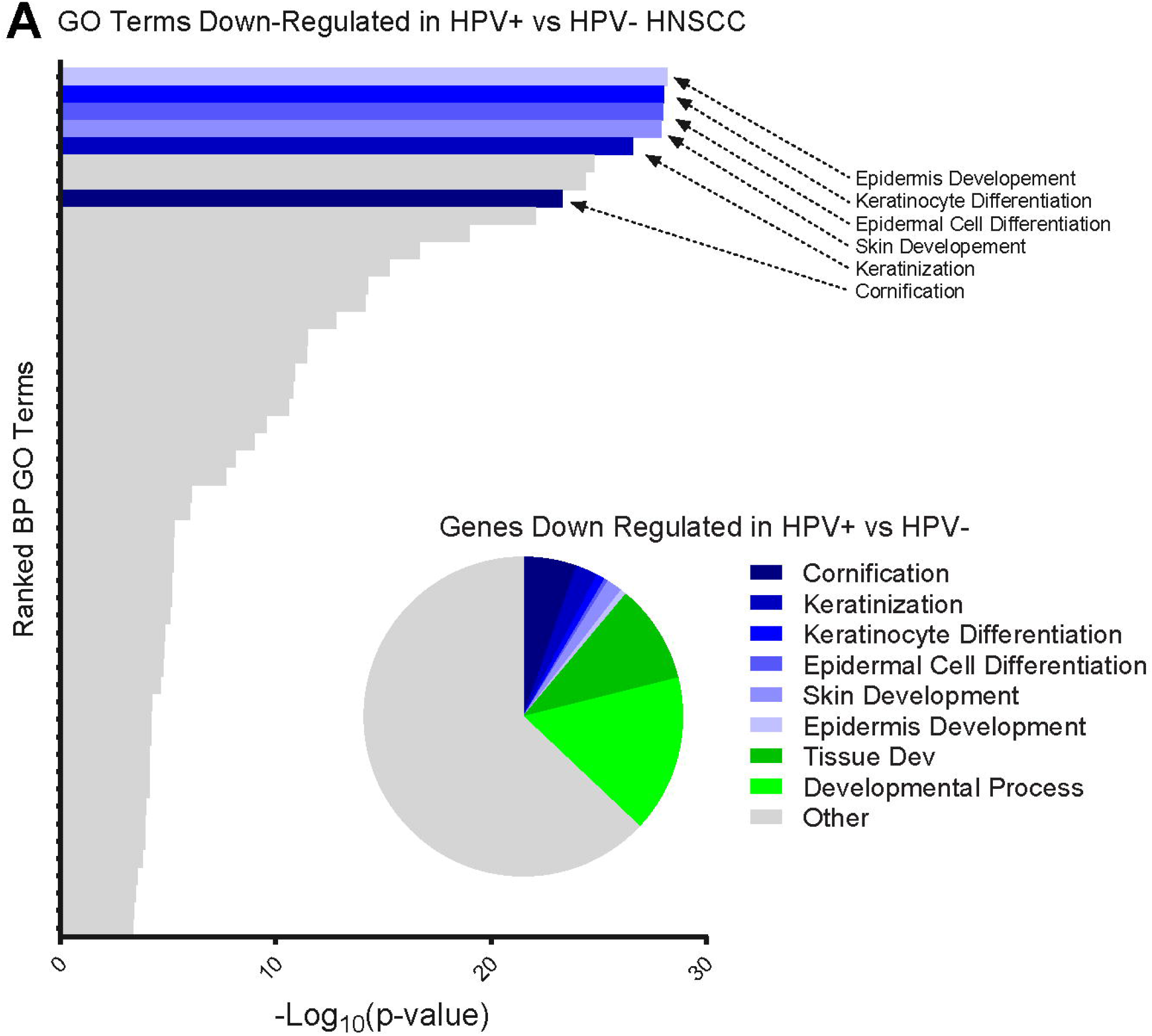
Keratinocyte differentiation gene signature describes the major differences between HPV+ and HPV-HNSCC. Data from HNSCC samples on the TCGA database were determined to be HPV+ or HPV- and analyzed for differences in gene expression. (A) Bar chart portrays the ranked –Log_10_(p-values) of enriched gene ontology terms among genes down-regulated in HPV+ HNSCC. Pie chart displays fraction of total down-regulated genes that fall into selected gene ontology categories.

## Discussion

Our previous finding that PTPN14 is targeted for degradation by high-risk HPV E7 but not by low-risk HPV E7 suggested that PTPN14 loss might be related to the biology of the high-risk HPV (47). PTPN14 is a candidate tumor suppressor based on the observation that it is mutated in some cancers (54, 57, 78–81). The targeted degradation of PTPN14 by high-risk HPV E7 requires the E3 ubiquitin ligase UBR4 and the interaction of UBR4 with papillomavirus E7 is required for E7 to transform cells (45, 46). Thus, PTPN14 degradation could be analogous to the well-established ability of high-risk HPV E6 but not low-risk HPV E6 to target p53 for proteasome-mediated degradation using the E3 ubiquitin ligase UBE3A (82, 83). However, neither our previous studies nor those from another group provided insight regarding the downstream effects of E7-mediated PTPN14 degradation in human keratinocytes (47, 84).

PTPN14 has been implicated as a negative regulator of YAP1, a transcriptional coactivator that is regulated by the Hippo signaling pathway (59, 61, 85). An appealing hypothesis was that E7-mediated PTPN14 degradation would activate YAP1 and promote the expression of pro-proliferative YAP target genes such as *CTGF* and *CYR61*. However, we have not identified any cell type in which high-risk HPV E7 expression causes an increase in *CTGF* or *CYR61* RNA. In addition, we found that depletion or knockout of PTPN14 in human keratinocytes did not cause *CTGF* or *CYR61* upregulation (Figures 4 and 5). However, our cell detachment experiments suggested that these genes are indeed regulated by Hippo signaling in keratinocytes (Figure 5). Thus, our results suggest that PTPN14 may not regulate Hippo-YAP signaling in keratinocytes.

In the absence of support for this initial hypothesis, we took an unbiased approach to determine the effect of high-risk HPV E7-mediated PTPN14 degradation in keratinocytes. By using an HPV16 E7 variant that cannot degrade PTPN14 (Figures 1 and 2) and by directly testing the effect of *PTPN14* knockout in primary HFK (Figure 4), we determined that PTPN14 loss results in a downregulation of several markers of epidermal cell differentiation. Consistent with this idea, *PTPN14* appears to be a target of regulation by p53 in mouse cells, but is likely a p63 target in human cells (79, 86–88). p63 is a master regulator of epidermal development (89). The link between PTPN14 and differentiation directly connected PTPN14 degradation to HPV biology.

To further test how high-risk HPV E7-mediated PTPN14 degradation affects processes related to epidermal cell differentiation, we used a keratinocyte detachment and re-plating assay (Figure 3). Our studies indicated that high-risk HPV E7 inhibit the expression of differentiation markers following cell detachment in a PTPN14 degradation-dependent manner. The same inhibition of differentiation markers occurred in detached PTPN14 knockout primary HFK (Figure 5). Anoikis is cell death triggered by detachment from a substrate and the ability to survive anoikis and proliferate in the absence of contact with the basement membrane is a hallmark of cancer cells. The E7 proteins that inhibited differentiation marker gene expression promoted cell survival following detachment and this correlated with the ability to degrade PTPN14 (Figure 3).

In support of the notion that E7 mediated PTPN14 degradation contributes to oncogenic transformation, our subsequent experiments indicated that PTPN14 degradation by high-risk HPV contributes to keratinocyte immortalization. Primary keratinocytes were fully immortalized by HPV16 E6/E7 but not by HPV16 E6/E7 E10K (Figure 6). In transcriptional profiles of human head and neck cancer samples, changes in gene expression consistent with PTPN14 loss were reflected in HPV-positive but not HPV-negative cancers (Figure 7). Strikingly, we found that the gene ontology terms related to keratinocyte differentiation and epidermis development described both the PTPN14-dependent differential gene expression in primary cells and the most significant differences between HPV-positive and HPV-negative head and neck carcinomas. We also observed that in previously published data these same GO terms were downregulated by the co-expression of HPV16 E6 and E7 in primary HFKs (90). These findings are consistent with the effect of PTPN14 loss being maintained throughout HPV-mediated carcinogenesis. Notably, the HPV16 E7 E10K variant that cannot bind UBR4 or degrade PTPN14 (Figure 3) was identified in a CIN3 lesion (62). We hypothesize that other patient-specific genetic or epigenetic changes may have compensated for the inability of E7 to degrade PTPN14 in this lesion, however only viral sequence information was collected from the patient samples in this study. Alternatively, this mutation may have impaired the progression of this lesion from CIN3 to a malignant cancer.

Some previous studies suggested that differentiation inhibition by E7 could be RB/E2F-dependent. E2F transcription factors have been shown to limit keratinocyte differentiation (91) and both the RB1-binding domain and the N-terminus of HPV16 E7 contributed to HPV-mediated differentiation inhibition in one study (20). However, our unbiased transcriptional analysis clearly showed that much of the E7-mediated repression of differentiation is independent of RB binding. Here we have focused on the genes repressed by HPV16 E7, which includes many markers of keratinocyte differentiation that were also downregulated upon PTPN14 knockout (Figures 2 and 4). Using the HPV16 E7 Δ21-24 mutant we have found that RB1 binding is not required for the repression of most of these genes (Figure 2). RB1 binding did allow for the repression of some differentiation genes by HPV16 E7 (Supplemental Figure 2D) and certain genes upregulated by HPV16 E7 but not by HPV16 E7 Δ21-24 were related to keratinocyte differentiation. Nonetheless, differentiation-related GO terms comprised a minor part of the HPV16 E7 gene induction signature whereas they were the most significant terms repressed by HPV16 E7 in the presence or absence of RB1 binding.

All HPVs, not only the high-risk types, likely manipulate differentiation in order to replicate. PTPN14 is a conserved interactor of HPV E7 suggesting an evolutionary pressure to maintain this interaction regardless of the ability to direct it for proteasomal degradation. Our future studies will address whether low-risk HPV E7 impact keratinocyte differentiation via their ability to bind to (but not degrade) PTPN14. Our study supports this hypothesis as the HPV16 E7 E10K variant and the low-risk HPV6 E7 proteins both bind PTPN14 without directing it for degradation and both inhibited keratinocyte differentiation to similar levels after detachment (Figure 5). More broadly, genus beta HPV E6 proteins bind MAML1 to inhibit Notch signaling, resulting in impaired keratinocyte differentiation and a cellular environment more conducive to virus replication (92–94). The genus alpha HPVs, which include all of the high-risk and low-risk HPV discussed here, do not engage MAML1 in the same way. It is interesting to speculate that all HPV promote proliferation but that pathogenesis is related to their ability to further impair differentiation: via the E7-PTPN14 interaction in the case of mucosal, genus alpha HPV and via the E6-MAML interaction in the case of cutaneous, genus beta HPV.

The binding and degradation of RB1 is a major component of high-risk HPV E7-mediated transformation. However, many observations have suggested that there must be RB1-independent contributions to E7-mediated transformation. We propose that PTPN14 degradation is a critical contributor to the oncogenic activity of high-risk HPV E7 and that PTPN14 inactivation impairs keratinocyte differentiation. PTPN14 degradation is conserved across the high-risk HPV E7 that we have tested so far and it is not dependent on the ability of E7 to bind or inactivate RB1. We have not yet established whether PTPN14 binding is sufficient to impair differentiation or whether PTPN14 degradation has additional oncogenic effects. In either case, the identification of a mechanism by which HPV E7 controls differentiation is significant. The potential of differentiation therapy has been validated by the highly successful use of all-trans retinoic acid to treat acute promyelocytic leukemia (95). It is tantalizing to speculate that inhibiting PTPN14 inactivation could similarly restore the cellular differentiation program in HPV-positive cancer cells and have therapeutic potential. Our future work will aim to elucidate the mechanism of PTPN14 signal transduction in keratinocytes and to further characterize the role of PTPN14 degradation in HPV replication and in HPV-associated cancers.

## Materials and Methods

### Cells

Primary human foreskin keratinocytes (HFK) (G5-Ep isolate, gift of James Rheinwald, Harvard Medical School) and hTert-immortalized HFK (63) were cultured as previously described (96). PTPN14 knockout or nontargeting control primary HFK were established by transduction with LentiCRISPR v2 vectors (Supplemental Table 4) followed by puromycin selection. N/Tert-Cas9 cells were generated by transduction with pXPR_111 (Addgene #59702) and blasticidin selection. PTPN14 knockout or mock control N/Tert cell lines were established by transfection of N/Tert-Cas9 with sgRNA targeting PTPN14 (Synthego, Supplemental Table 4). Retroviruses and lentiviruses were generated as previously described (96).

To assess keratinocyte survival and changes in gene expression following detachment from a substrate, N/Tert-mock or N/Tert-sgPTPN14 cells were harvested by trypsinization and re-plated in ultra-low attachment plates (Sigma-Aldrich CLS3471). After 0 or 12h of culture in suspension cells were harvested for RNA analysis or 1000 cells were re-plated in standard 6-well tissue culture plates. Re-plated cells were stained with crystal violet 5 days post re-plating.

To assess the ability of HPV16 E7 variants to support keratinocyte immortalization, primary HFK were transduced with one MSCV-based retroviral vector encoding conferring puromycin resistance (HPV16 or HPV6 E6 or an empty vector control) and one retroviral vector conferring blasticidin resistance (HPV16 or HPV6 E7 or an empty vector control) (Supplemental Table 4). Cells were selected in puromycin and blasticidin and passaged for approximately 110 days. Population doublings were calculated based upon the number of cells collected and re-plated at each passage.

### Plasmids and cloning

LentiCRISPR v2 vectors were cloned according to standard protocols using sgRNA sequences as contained in the Broad Institute Brunello library (97). The E10K mutation was introduced by site-directed mutagenesis into pDONR-Kozak-16E7 and recombined into MSCV-IP N-FlagHA GAW as previously described (96). Additional HPV E6 and E7 retroviral vectors used in the study are listed in Supplemental Table 4.

### Western blotting

Western blots were performed as previously described (47) using Mini-PROTEAN or Criterion (BioRad) SDS-PAGE gels and transfer to PVDF. Membranes were blocked in 5% nonfat dried milk in TBS-T (Tris buffered saline [pH 7.4] with 0.05% Tween-20), then incubated with primary antibodies as follows: RB1 (Calbiochem/EMD), actin (Millipore), PTPN14 (R&D Systems), and UBR4 (gift of Dr. Yoshihiro Nakatani, Dana-Farber Cancer Institute (98)). Membranes were washed in TBS-T and incubated with horseradish peroxidase (HRP)-coupled anti-mouse or anti-rabbit antibodies or an Alexa-680 coupled anti-mouse antibody and detected using Western Lightning chemiluminescent substrate or a Li-COR Infrared imaging system. HA-tagged proteins were detected using an HA antibody conjugated to HRP (Roche) and visualized in the same way. For anti-HA immunoprecipitations, HA-tagged proteins were immunoprecipitated and processed for Western blot as previously described (47).

### RNA-seq

Total RNA was isolated from 2-3 independent isolates of HFK-control, HFK-PTPN14 KO, HFK-empty vector control, or HFK E7 cells using the RNeasy mini kit (Qiagen). PolyA selection, reverse transcription, library construction, sequencing, and initial analysis were performed by Novogene. Differentially expressed genes were selected based on a 1.5-fold change and adjusted *p*≤0.05 cutoff and were analyzed for enriched biological processes (BP) using the GO enrichment analysis tool of the PANTHER classification system (99). All GO terms in enrichment analyses are displayed in rank order by adjusted *p*-value. RNA-seq data have been deposited in NCBI GEO with accession number GSE121906.

### qRT-PCR

Total RNA was isolated from N/Tert cells using the NucleoSpin RNA extraction kit (Macherey-Nagel). RNA was then reverse transcribed using the High Capacity cDNA Reverse Transcription Kit (Applied Biosystems). cDNAs were assayed by qPCR using Fast SYBR Green Master Mix (Applied Biosystems) using a QuantStudio 3 - 96-Well, 0.2 mL Block instrument (ThermoFisher). All gene RT-qPCR data were normalized to GAPDH or to G6PD. qRT-PCR primer sequences are listed in Supplemental Table 4.

## Acknowledgements

We thank the members of our laboratories for helpful discussions and suggestions and Yoshihiro Nakatani for the anti-UBR4 antibody. This work was supported by American Cancer Society grant 131661-RSG-18-048-01-MPC to E.A.W. and by National Institutes of Health R01 CA066980 to K.M.

## Author Contributions

Conceptualization, J.H., K.M., and E.A.W.; Formal Analysis, J.H., T.J.N., B.W., I.M.M., and E.A.W.; Investigation, J.H., A.E.B., M.G., and E.A.W.; Resources, T.J.N., B.W., I.M.M., M.G., and K.M.; Data Curation, J.H. and E.A.W.; Writing – Original Draft, J.H. and E.A.W.; Writing – Review and Editing, J.H., I.M.M., K.M., and E.A.W.; Visualization, J.H. and E.A.W.; Supervision, E.A.W.

## Declaration of Interests

The authors declare no competing interests.

## Supplemental Figure and Table Legends

**Supplemental Figure 1. The HPV16 E10K variant is impaired in PTPN14 degradation but binds RB1 and promotes cell cycle progression.**

Primary HFK were transduced with retroviruses encoding HPV16 E7 WT, HPV16 E7 E10K, HPV16 E7 Δ21-24, or an empty vector control. RNA from transduced cells was polyA selected and analyzed by RNA-seq for changes in gene expression. (A) Heat map displays top 75 genes differentially expressed in HPV16 E7Δ21-24 relative to HPV16 E7 WT cells. (B) Genes in the DNA Replication GO Term (GO0006260) altered by HPV16 E7 WT relative to control ≥1.5 fold with *p*-value ≤0.05 are displayed in heat map.

**Supplemental Figure 2. HPV 16E7 degrades PTPN14 to inhibit keratinocyte differentiation.**

Primary HFK were transduced with retroviruses encoding HPV16 E7 WT, HPV16 E7 E10K, HPV16 E7 Δ21-24, or an empty vector control. RNA from transduced cells was polyA selected and analyzed by RNA-seq for changes in gene expression. (A) GO enrichment analysis of genes >1.5 fold higher with p<0.05 in HPV16 E7 WT relative to HPV 16 E7 E10K shows no strong enrichment for GO terms. (B) Same analysis as (A) of (C, Top) genes with ≥1.5 fold lower expression in HPV16 E7 WT than HPV16 E7 Δ21-24 cells, and (C, Bottom) genes ≥1.5 fold higher expression in HPV16 E7 Δ21-24 cells relative to empty vector control cells and *p*-value ≤0.05. (C) Scatter plot of log2(fold-change) in gene expression compares the gene expression changes of HPV16 E7 Δ21-24 relative to empty vector control to those of PTPN14 KO relative to control. Colors denote whether genes are altered by PTPN14 KO only (blue), by HPV16 E7 Δ21-24 only (light green), or both (dark green). (D) Unbiased clustering of genes that are lower in HPV16 E7 WT cells relative to HPV16 E7 E10K by ≥1.5 fold with *p*-value ≤0.05 in HPV16 E7. Gene names and clustering are displayed to the left of the heat map, and selected GO categories are displayed on the right. Color coding on the right side denotes whether genes in a cluster are related to epidermis development (blue), other developmental processes (green), or neither (gray).

**Supplemental Figure 3: PTPN14 depletion upregulates inflammatory response genes in primary human keratinocytes and down regulates similar keratinocyte differentiation genes to HPV16 E7.**

Primary HFK were transduced with LentiCRISPRv2 lentiviral vectors encoding SpCas9 and non-targeting or PTPN14-directed sgRNAs and analyzed for changes in gene expression. PolyA selected RNA was analyzed by RNA-seq. Plot displays GO enrichment analysis of genes upregulated in HFK-PTPN14 KO compared to HFK-control.

**Supplemental Figure 4. Keratinocyte differentiation gene signature describes the major differences between HPV+ and HPV-HNSCC.**

Data from HNSCC samples on the TCGA database were determined to be HPV+ or HPV- and analyzed for differences in gene expression. (A) Bar chart portrays the ranked –Log_10_(p-values) of enriched gene ontology terms among genes up-regulated in HPV+ HNSCC.

**Supplemental Table 1. The HPV16 E10K variant is impaired in PTPN14 degradation but binds RB1 and promotes the expression of E2F-regulated genes.**

Primary HFK were transduced with retroviruses encoding HPV16 E7 WT, HPV16 E7 E10K, HPV16 E7 Δ21-24, or an empty vector control. RNA from transduced cells was polyA selected and analyzed by RNA-seq for changes in gene expression. (A) Table includes top 75 genes significantly altered by HPV16 E7 Δ21-24 relative to HPV16 E7 WT. (B) Table displays the genes in the DNA Replication GO Term (GO0006260) that are altered by ≥1.5 fold with p-value ≤0.05 in HPV16 E7 WT cells relative to vector control cells. Tables include gene name, log2(fold change), and adjusted p-value for HPV16 E7 E10K and HPV16 E7 Δ21-24 relative to HPV 16 E7 WT.

**Supplemental Table 2. HPV 16E7 degrades PTPN14 to inhibit keratinocyte differentiation.**

Primary HFK were transduced with retroviruses encoding HPV16 E7 WT, HPV16 E7 E10K, HPV16 E7 Δ21-24, or an empty vector control. RNA from transduced cells was polyA selected and analyzed by RNA-seq. (A) Table includes gene name, log2(fold change), and adjusted p-value for genes altered by ≥1.5 fold with *p*-value ≤0.05 in HPV16 E7 WT relative to HPV16 E7 E10K. (B) Table includes gene name, log2(fold change), and adjusted p-value for genes altered by ≥1.5 fold with *p*-value ≤0.05 in HPV16 E7 Δ21-24 relative to empty vector control.

**Supplemental Table 3: PTPN14 depletion impairs differentiation-related gene expression in primary human keratinocytes.**

Primary HFK were transduced with LentiCRISPRv2 lentiviral vectors encoding SpCas9 and non-targeting or PTPN14-directed sgRNAs and polyA selected RNA was analyzed by RNA-seq. Table includes gene name, log_2_(fold change), and adjusted p-value for genes differentially expressed by ≥1.5 fold with p-value ≤0.05.

**Supplemental Table 4: Plasmids and primers used in the study.**

